# A switch in collagen expression regulates cessation of distal tip cell migration in *C. elegans*

**DOI:** 10.1101/2025.10.03.680101

**Authors:** Victor Stolzenbach, Sophie S. Griffin, Priti Agarwal, Ronen Zaidel-Bar, Erin J. Cram

**Affiliations:** Department of Biology, Northeastern University, Boston, MA, USA; Department of Cellular, Developmental, and Regenerative Biology, Gray School of Medical Sciences, Gray Faculty of Medical and Health Sciences, Tel Aviv University, Tel Aviv, Israel

**Keywords:** Cell invasion, leader cell, collagen, gonad morphogenesis, *C. elegans*

## Abstract

Development of the *C. elegans* gonad requires precise regulation of cell migration. The distal tip cell (DTC) guides the elongation of the gonad into its final U-shaped structure before halting in adulthood. How cessation of elongation is regulated remains unknown. Here, we analyze an RNA-seq data set isolated from stage-specific DTCs to uncover the temporal gene expression dynamics underlying this process. Collagens emerged as the most enriched gene family during the transition from migratory larval stages to adulthood. We identify distinct temporal waves of collagen expression, culminating in a core adult-specific module that coincides with migration cessation. Functional analysis by RNAi depletion revealed that many collagens upregulated in adulthood are required for timely migration arrest, while others affect gonad shape. Perturbation of collagen remodeling enzymes phenocopied these effects. Our findings uncover a stage-specific collagen program in the DTC and suggest that terminal migration arrest is actively reinforced by matrix remodeling.

## Introduction

Cell migration is fundamental to organogenesis and tissue patterning across species. Equally important as initiation and maintenance of cell migration is the timely cessation of migration, a critical step required for correct development, tissue homeostasis, and to prevent metastatic disease progression (Bravo-Cordero et al., 2012). For instance, in developmental contexts such as neural crest migration, precise control over migration termination is crucial to avoid developmental abnormalities (Bronner and Simões-Costa, 2016). Yet, many questions remain about the how cells know when and where to stop migrating.

The hermaphrodite gonad of *Caenorhabditis elegans* provides a well-established model to study this process. The formation of the gonad is guided by specialized leader cells called distal tip cells (DTCs), which migrate throughout the larval stages and cease migration at adulthood. The DTCs initially navigate along the ventral body wall muscle during the second larval stage (L2), turn towards the dorsal side of the animal during the third larval stage (L3), and finally guide elongation along the dorsal body wall in the fourth larval stage (L4) until they cease movement in early adulthood (Agarwal et al., 2022; Blelloch and Kimble, 1999). This migration is propelled by the proliferative pressure from the germline behind the DTC, which pushes these leader cells forward.

Directional steering and turning are achieved through polarized cell-matrix adhesion, particularly via integrin-mediated interactions (Agarwal et al., 2022). The DTC locally secretes matrix metalloproteases (MMPs), such as GON-1/ADAMTS, to degrade the extracellular matrix (ECM), facilitating forward progression by relieving physical resistance at the migration front (Agarwal et al., 2022; Blelloch and Kimble, 1999; Nishiwaki et al., 2000). At the end of larval morphogenesis, each DTC comes to a stop at the midline of the animal and remains stationary during adulthood. Large-scale genetic screens have identified extensive gene networks regulating various aspects of this migration process, emphasizing the complexity and coordination required for proper DTC guidance (Cram et al., 2006).

Several regulators of the termination of DTC migration and gonad elongation have been identified, including the conserved transcription factor VAB-3/Pax6. In *vab-3* mutant animals, DTC migration continues in adulthood, resulting in elongated and coiled ‘cinnamon roll’ shaped gonad arms (Nishiwaki, 1999; Zhang and Emmons, 1995). VAB-3 induces an integrin switch, reducing the expression of INA-1 α integrin and promoting the expression of PAT-2 α integrin, thereby presumably altering cell-matrix interactions necessary for migration termination (Meighan and Schwarzbauer, 2007). The spliceosome regulators, CACN-1/Cactin and CCDC-55, are also required for the DTC to stop migrating at the end of L4. Depletion of either of these factors by RNAi leads to migration of the DTC well past the midpoint of the animal (Kovacevic et al., 2012; Tannoury et al., 2010). These results suggest there may be a switch in the DTC transcriptome at the end of L4 that regulates the cessation of DTC migration.

Collagens influence morphogenesis by providing structure, through biochemical signaling interactions, and by regulating tissue mechanical properties such as stiffness (Page-McCaw et al., 2007; Rozario and DeSimone, 2010). *C. elegans* encodes over 170 collagen genes, with the majority classified as cuticle collagens involved in molting and external structure. Unlike vertebrate fibrillar collagens, which assemble into long, rope-like fibers, *C. elegans* cuticle collagens are cross-linked through tyrosine-derived di- and tri-tyrosine bonds that provide tensile strength and flexibility. This specialized cuticle matrix is synthesized anew at each molt to form the collagen-based exoskeleton (Page and Johnstone, 2007). However, only a small subset of these collagens have been functionally characterized, and their roles in internal tissues remain largely unexplored (Kramer, 1994). The basement membrane type IV collagens EMB-9 and LET-2 form critical scaffolds essential for accurate DTC navigation, and their disruption leads to severe migration defects, including wandering or stalled DTCs (Gupta et al., 1997; Kubota et al., 2008). The metalloprotease MIG-17, secreted by body wall muscle, specifically remodels type IV collagen to ensure its proper distribution along the gonadal basement membrane. Its loss results in abnormal collagen accumulation, impaired nidogen recruitment, defective basement membrane assembly, and disrupted DTC migration, ultimately compromising gonad shape and healthspan (Kubota et al., 2004; Kubota et al., 2008; Nishiwaki et al., 2000; Shibata et al., 2024). CLE-1/type XVIII collagen, which contains an endostatin-like domain, promotes distal tip cell migration via its NC1 domain. Loss of *cle-1* leads to premature dorsal turns and misdirected gonad arm outgrowth, indicating that CLE-1 provides promigratory cues required for proper DTC navigation (Ackley et al., 2001; Gupta et al., 1997; Jayadev et al., 2019). In mammalian systems, collagen XVIII regulates endothelial cell migration and angiogenesis, and disruptions are associated with pathological conditions such as fibrosis, tumor progression, and angiogenesis (Bonnans et al., 2014; Marneros and Olsen, 2005).

By analyzing a stage-specific RNAseq data set derived from larval and adult DTCs, we discovered that numerous collagens display stage-specific expression, suggesting functional specialization during migration and stopping (Agarwal et al., 2025). Targeting these collagens by RNAi-mediated knockdown demonstrated that many of them are required for migration cessation, with depletion leading to continued DTC movement beyond the normal stopping point. Additionally, RNAi knockdown of key DTC-expressed collagens caused gonad deformation, underscoring their essential role in maintaining gonad integrity.

## Results

### Transcriptomic Shifts in the Extracellular Matrix Highlight Collagen Dynamics in the Distal Tip Cell

The DTC guides elongation of the U-shaped hermaphrodite gonad, navigating two turns before coming to a stop at the animal’s midline. The gonad arms are surrounded by a collagen-rich extracellular matrix. The type IV collagen α2 ortholog LET-2::mNG outlines the gonad (Figure 1A). Previous work has described the role of LET-2 and many other regulators in DTC migration, however, much less is known about the factors that lead to cessation of gonad elongation in adults (Kubota et al., 2008).

**Figure 1.**
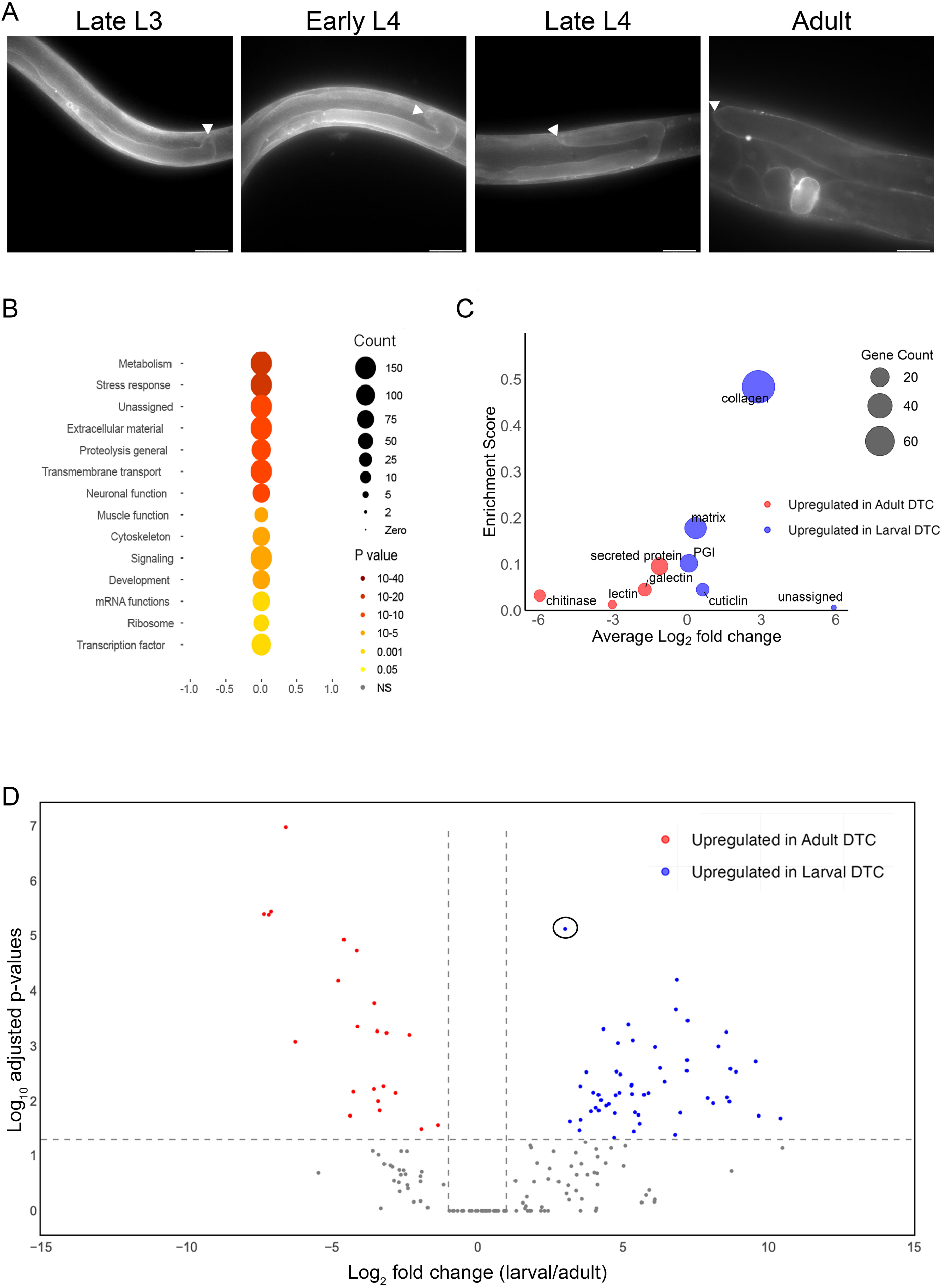
Collagen category enrichment and stage-specific expression in distal tip cells. **(A)** Representative images of the *C. elegans* gonad expressing LET-2::mNG at late L3, early L4, late L4, and adult stages. mNeonGreen signal outlines the gonad, highlighting collagen localization along the basement membrane during and after DTC migration (arrows denote DTC). **(B)** WormCat analysis of differentially expressed genes (DEGs) in larval vs. adult DTCs. Bubble size indicates gene count; shading intensity reflects statistical significance. The p-value scale spans from 10^−^ □ □ (darkest, strongest enrichment) to 0.05 (lightest, weakest enrichment); NS indicates non-significant categories. Categories such as metabolism, stress response, and extracellular material were most significantly altered. **(C)** Subcategory-level visualization of ECM-related gene expression from the “Extracellular Material” WormCat category. Each bubble represents a subcategory, with size proportional to the number of genes, y-axis showing enrichment score, and x-axis showing average log_2_ fold change (larval vs. adult). Blue bubbles indicate larval enrichment; red bubbles indicate adult enrichment. **(D)** Volcano plot of collagen gene expression differences between larval and adult DTCs. Collagens upregulated in larval DTCs are marked in blue; those upregulated in adults are in red. One gene, *let-2*, is circled. Vertical dashed lines mark ±1 log_2_ fold change, and the horizontal dashed line indicates the significance threshold (adjusted p = 0.05; log_10_ value = 1.3).

To identify genes involved in DTC migration, GFP-tagged DTCs were isolated from *C. elegans* at late L3, early L4, late L4, and young adult stages, along with matched non-GFP cells as controls. Bioinformatic analyses were performed to comprehensively characterize developmental stage-specific transcriptional profiles, as described previously (Agarwal et al., 2025). First, we identified and characterized the genes differentially expressed between larval and adult DTCs, as a strategy to identify genes required for cessation of DTC migration. Among all significantly regulated categories, genes associated with extracellular material, defined as genes linked to formation of the extracellular matrix, cuticle, or secreted proteins, showed the highest degree of enrichment, marked by both gene count and statistical significance (Figure 1B). Collagens represented the most enriched subgroup within the extracellular material category (Figure 1C). Other prominent categories included metabolism, stress response, proteolysis, and transmembrane transport.

To further explore temporal trends in ECM-related gene expression, we compared larval and adult DTCs across extracellular matrix subcategories (Figure 1C). Many collagen-related categories appeared more enriched in larval stages, suggesting that migrating DTCs are actively remodeling their environment. However, we noticed that a subset of these larval-enriched collagens remain highly expressed in adult DTCs (Figure 2B). This raised the possibility that these collagens play continued roles in adulthood, prompting us to more systematically investigate collagen gene dynamics across developmental stages.

**Figure 2.**
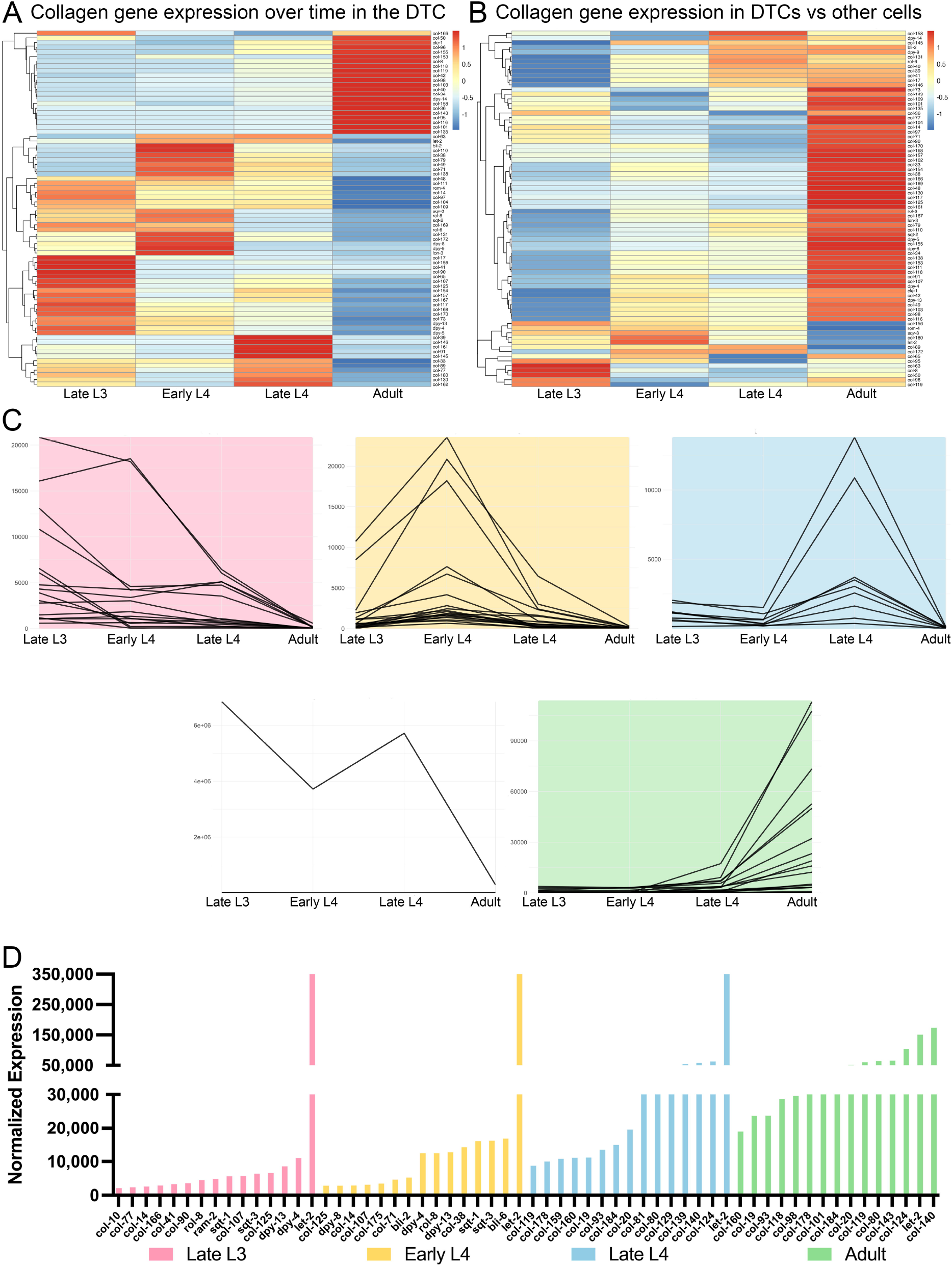
Collagen gene expression dynamics and temporal clustering across DTC development. (A) Heatmap showing z-scored expression of significantly expressed collagens in the DTC across four developmental stages. Genes were clustered by expression pattern. (B) Heatmap of the same collagens, comparing DTC expression to all other cells (somatic and germline) at each stage based on scaled log_2_ fold change values. (C) Mfuzz clustering reveals five dominant temporal expression trajectories among differentially expressed collagens, highlighting stage-specific activation patterns. The unshaded panel at lower left corresponds to the cluster containing *let-2*, which displays a biphasic pattern, high in late L3, down in early L4 stages, then rising again in late L4 before declining in adulthood, dominating the cluster and obscuring the other 11 genes. (D) Bar graph of normalized expression of the highest-expressed collagen genes at each stage. The top 15 expressed collagens per stage are shown. A split y-axis is used to improve visualization: the first segment increments by 10,000 up to 30,000, then jumps to 50,000 and increases by 100,000 thereafter. Expression values are TMM-normalized CPM averaged across replicates.

**Figure 3.**
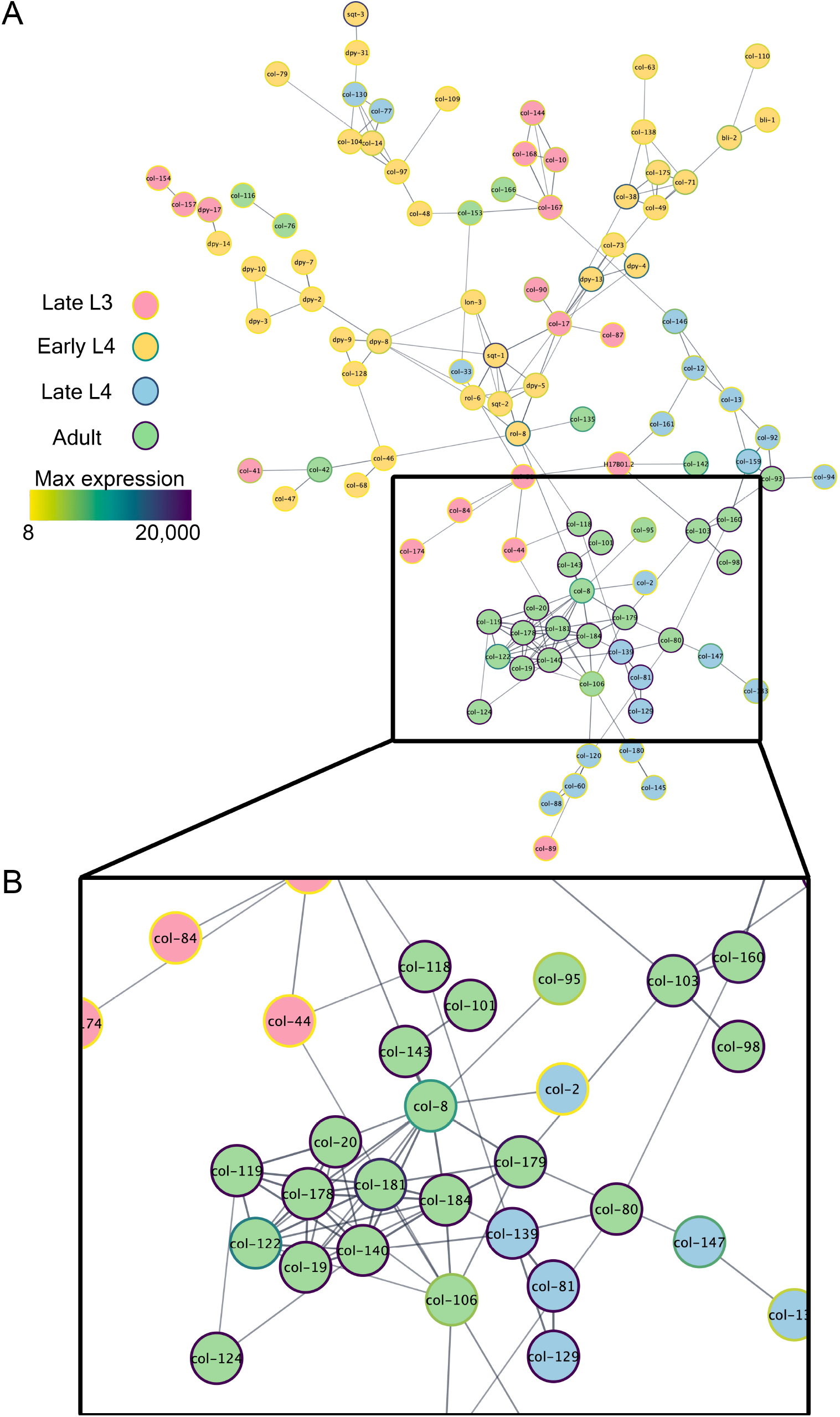
Collagen interaction network highlights stage-specific coordination during DTC development. (A) Collagen interaction network generated using STRING-db at a confidence cutoff of 0.7. Each node represents a collagen gene and is colored by its stage of peak expression: pink (late L3), yellow (early L4), blue (late L4), and green (adult). The outline of each node is scaled by the gene’s maximum expression value, with purple representing high expression and yellow representing low expression. Edges represent known or predicted interactions, with thicker lines indicating stronger confidence scores. (B) Close-up of the adult collagen-enriched module, showing tightly connected collagens with high expression in adulthood. These genes cluster together, suggesting coordinated regulation and shared roles in post-migratory gonad structure.

We next curated a list of 184 genes annotated as collagens in the *C. elegans* genome. These genes include all members assigned by WormCat to collagen subcategories under “extracellular material,” ensuring broad coverage of the family. We then analyzed their expression dynamics across developmental stages using edgeR. This targeted analysis revealed strong temporal regulation, with 53 collagens significantly upregulated in larval DTCs and 23 enriched in adults (Figure 1D). Although expression of many collagen genes decreases in adult DTCs relative to larval stages, most genes remain enriched in DTCs compared to non-GFP somatic cells, underscoring their continued tissue-specific role.

To place these transcriptional patterns in an evolutionary context, we identified human orthologs of collagen genes affecting DTC migration based on sequence similarity. The analysis revealed that affected *C. elegans* collagens share homology with four major human collagen groups: type VI (COL6A5), type XXI (COL21A1), type IV (COL4A2), and type XXIV (COL24A1) (Supplemental Table 2). These clades correspond to collagen subgroups with known functions in cuticle integrity, vascular and neural structure, basement membrane formation, and skeletal development, respectively.

### Temporal Clustering Reveals Stage-Specific Collagen Expression Programs in Distal Tip Cells

To identify collagen genes that might play a role in migration and anchoring, we analyzed the expression of the same set of 184 collagen genes described earlier across developmental stages. Collagen gene expression exhibited dynamic changes from late L3 through adulthood, with stage-specific expression patterns revealed by z-scored heatmaps (Figure 2A). In this heatmap, expression values were normalized using z-scores, which reflect how strongly each gene is up- or down-regulated at a given stage relative to its own mean expression across all stages. This approach highlights temporal expression trends within each gene, allowing better visualization of stage-specific regulation. A comparison of collagen expression in DTCs versus other cells across matched timepoints revealed that many of these collagens showed a relative shift in expression during the adult stage (Figure 2B). Although many collagens peak during larval stages, many continue to be expressed significantly in adulthood, indicating sustained functional roles beyond development.

To further explore these patterns, we applied Mfuzz clustering to significantly differentially expressed collagens. Five distinct expression clusters were identified, each displaying a characteristic temporal trajectory (Figure 2C). The clustering analysis captured peak expression across developmental stages, suggesting coordinated transcriptional changes over time.

One cluster of eleven genes, dominated by *let-2*, showed a unique biphasic pattern: high in late L3, reduced during the intermediate stages, and reactivated in adults. This cluster’s profile, distinct from the others, may reflect a specialized role for *let-2* in both early migration and later stabilization, consistent with its previously observed function in maintaining gonad architecture (Kawano et al., 2009; Kubota et al., 2008). Each of the remaining four clusters was composed of collagens whose expression peaked at a specific developmental stage: late L3, early L4, late L4, or adult. In the late L3 cluster, expression levels declined gradually over time, whereas in the early and late L4 clusters, expression sharply increased at the peak stage and then decreased by adulthood.

Notably, expression of a subset of collagens, including *col-140, col-124, col-80, col-184*, begins in late L4 and continues into adulthood. To visualize overall expression magnitude, normalized read counts of the top 15 most highly expressed collagens at each stage were plotted (Figure 2E), revealing a progressive increase in collagen expression across development, with the adult DTC expressing the highest overall levels of collagen.

### Collagen and Collagen Remodeling Genes Are Required for Proper Distal Tip Cell Migration

To test the functional requirement for the collagens identified in our transcriptomic and clustering analyses, we individually depleted 37 collagen genes using DTC-specific RNAi (Supplementary Table 1 for all knockdowns). We prioritized the collagens shown in Figure 2E, specifically those with significant differential expression, and those with very high expression in late L4 and young adult DTCs, when migration is stopping. Beyond that, we included collagens with an adjusted p-value < 0.05 and an absolute log_2_ fold change > 1 between larval and adult stages. Among those, all collagens with available RNAi clones were tested. RNAi knockdown of collagen genes was restricted to the DTC using an *rde-1*(*ne219*);*rrf-3*(*pk1426*) strain in which the argonaut RDE-1 is expressed in the DTC, restoring sensitive RNAi activity specifically in DTCs. This allowed us to isolate cell-autonomous effects on migration and morphology (Figure 4A). Of these, 22 knockdowns caused significant DTC migration defects (Figure 4B). Collagens highly expressed in the adult DTC, such as *col-101, col-119, col-143, col-184*, and *col-19* produced significant overshoot phenotypes. Depletion of *col-178* caused the DTC to execute an extra turn, suggesting COL-178 is also required for directional control.

**Figure 4.**
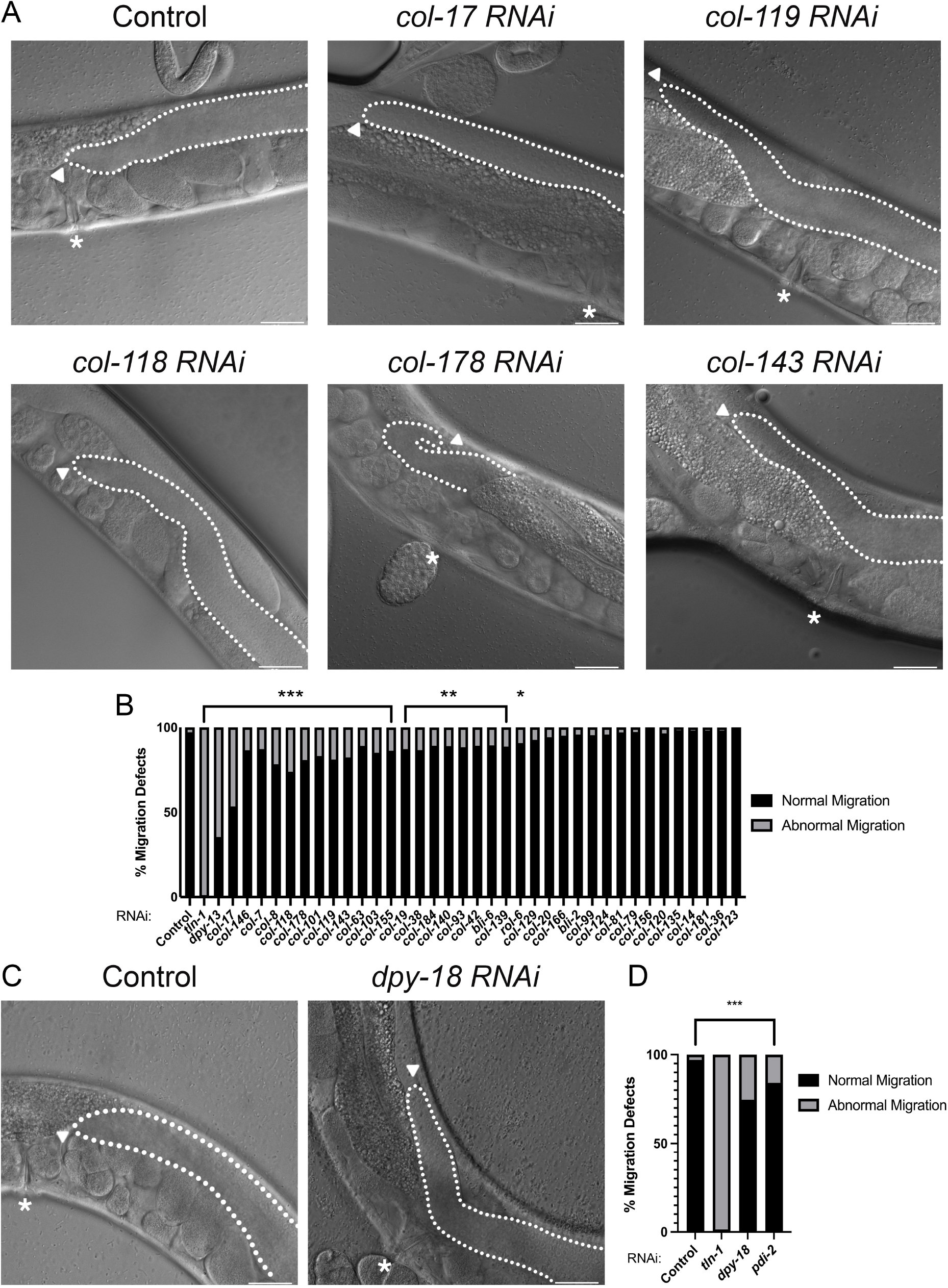
RNAi screen reveals collagen-dependent regulation of DTC migration. (A) Representative DIC images showing distal gonad arms in control worms and RNAi-treated animals targeting individual collagens. The gonad is outlined in white. Arrowheads denote the position of the distal tip cell (DTC), and asterisks indicate the vulva when visible. Several collagen knockdowns, including *col-17, col-119, col-118, col-178*, and *col-143*, exhibit aberrant DTC migration and gonad morphology. (B) Quantification of DTC migration defects across RNAi treatments. Bars indicate percentage of animals with normal (black) or abnormal (gray) migration. Fisher’s exact test with Benjamini-Hochberg correction was used to determine statistical significance (*p < 0.05, **p < 0.01, ***p < 0.001). (C) Representative images comparing control animals to worms treated with RNAi targeting *dpy-18* and *pdi-2*, both of which are genes involved in collagen remodeling. The gonad is outlined in white, arrowheads mark the DTC, and asterisks indicate the vulva when visible. RNAi knockdown of *dpy-18* resulted in penetrant DTC migration defects and abnormal gonad shaping. (D) Quantification of DTC migration defects following *dpy-18* and *pdi-2* RNAi, confirming that knockdown of collagen remodelers significantly disrupts DTC migration (*p < 0.05, **p < 0.01, ***p < 0.001). Scale bars in all images (panels A and C) represent 20 µm.

We next evaluated the role of collagen remodeling enzymes by targeting DPY-18 (P4HA1/2) and PDI-2 (P4HB), orthologs of human procollagen processing factors. Although these genes did not emerge from our transcriptomic analysis, we selected them based on their essential roles in cuticle collagen biogenesis, reasoning that they might similarly affect collagen remodeling in the DTC. DTC-specific RNAi against *dpy-18* caused severe migration defects and gonad deformation, while *pdi-2* knockdown led to milder defects (Figures 4C–D). Notably, *pdi-2* RNAi also produced germ cell disorganization and enlargement of the gonad arm, despite the RNAi being confined to the DTC, suggesting that collagen remodeling by the DTC may exert non-cell-autonomous effects on neighboring tissues.

In parallel, we observed that several collagen knockdowns resulted in abnormal gonad shapes, including expanded or bulbous distal arms (Figure 5A). RNAi targeting *bli-2, rol-6, col-93*, and *col-156* caused significant increases in gonad width relative to controls (Figure 5B), implicating these collagens in maintaining the structure of the gonad arm. Together, these findings demonstrate that collagen pathways are essential for DTC migration and gonad architecture, likely through both cell adhesion signaling and ECM-mediated structural support. Our working model proposes that the DTC is encased within a network of collagens that physically constrains its size and provides structural resistance during migration. Loss of collagen may disrupt this interconnected matrix, weakening the overall scaffold and preventing proper compaction, which can lead to overexpansion and failure to terminate migration appropriately (Figure 6).

**Figure 5.**
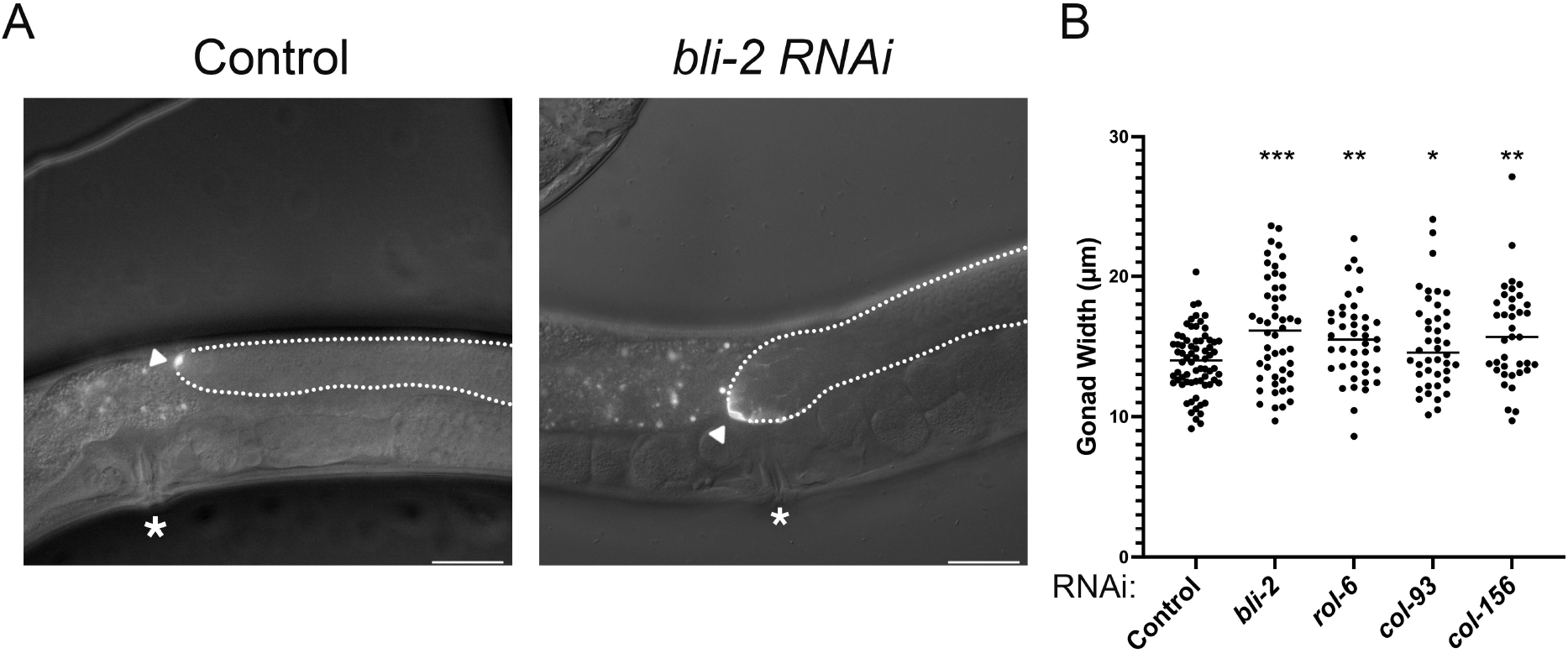
Collagen knockdown causes gonad widening defects in distal gonad arms. (A) Representative fluorescence micrographs showing gonad morphology in control and *bli-2* RNAi-treated animals using *lag-2p::mNG::PH* to label the distal tip cell. Knockdown of *bli-2* leads to a visibly expanded gonad. Scale bars, 20 µm. (B) Quantification of gonad width (µm) following RNAi treatment targeting *bli-2, rol-6, col-93, and col-155*. All four genes caused a significant increase in gonad width compared to controls (Welch’s t-test, *p < 0.05, **p < 0.01, ***p < 0.001).

**Figure 6.**
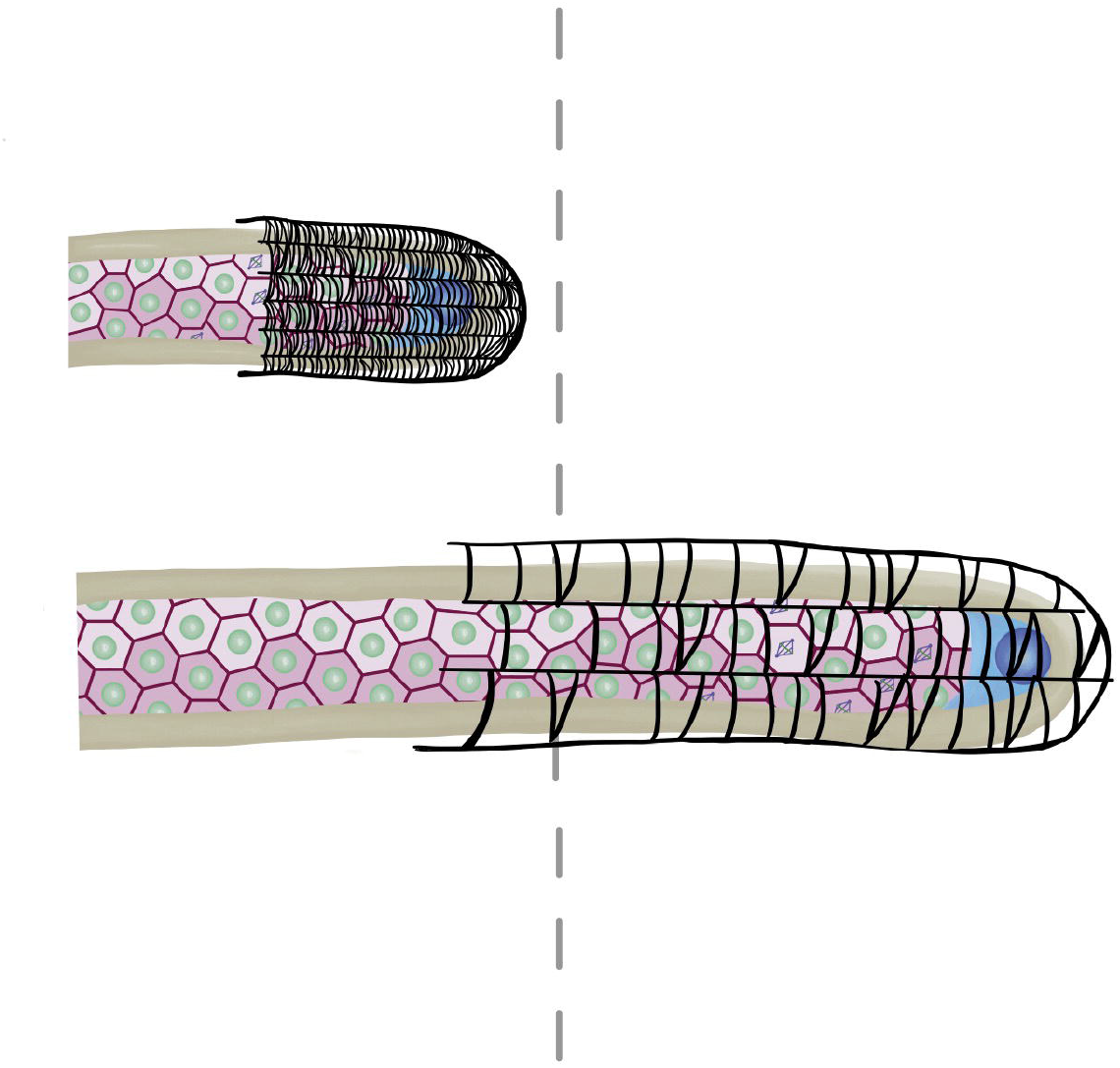
Model for collagen-dependent regulation of distal tip cell migration cessation. Schematic illustrating a proposed model for how collagens regulate the timing of distal tip cell (DTC) migration arrest. (Top) In wild-type animals, adult-enriched collagens assemble into a structured extracellular matrix “net” surrounding the distal gonad arm. This network progressively constrains DTC movement, providing mechanical resistance that facilitates timely cessation at the midline. (Bottom) In collagen-depleted animals, the network is weakened, akin to missing links in a chain. Reduced matrix integrity delays migration arrest, allowing the DTC to overshoot its normal stopping point and producing gonad defects.

## Discussion

Our study reveals that distal tip cell (DTC) migration in *C. elegans* is associated with a distinct stage-specific transcriptional program that includes collagen expression. A defined set of collagens becomes upregulated as DTCs halt their movement in adulthood, suggesting that a specific subset of collagen genes regulates migration cessation. RNAi-mediated depletion of these adult-enriched collagens leads to persistent migration and disrupted gonad morphology, indicating that they are required for cessation of gonad elongation. One possibility is that the DTC deposits ECM that constrains its own movement and final positioning. These findings align with a growing body of work reframing the ECM as an active driver of morphogenesis rather than a passive scaffold. In other systems, patterned ECM remodeling can generate forces, alter tissue stiffness, and guide shape changes independently of cellular contractility (Díaz-de-la-Loza and Stramer, 2024). For example, anisotropic basement membrane deposition in the *Drosophila* egg chamber biases organ elongation, and collagen network assembly in the ventral nerve cord produces intrinsic stresses that drive condensation (Crest et al., 2017; Haigo and Bilder, 2011; Matsubayashi et al., 2017; Serna-Morales et al., 2023). By analogy, our identification of a stage-specific collagen program in the DTC suggests that targeted collagen deposition and remodeling may directly create a mechanical or structural barrier that halts migration.

Our results support a model in which adult-enriched collagens reinforce migration arrest. For instance, depletion of *col-119* and *col-143* by RNAi allows the DTC to continue migrating into adulthood. This contrasts with CLE-1/collagen XVIII, whose NC1 domain promotes neuronal and axon migration, and whose loss leads to premature termination or misrouting (Ackley et al., 2001). Similar roles for type IV collagen in guiding epithelial morphogenesis have been described in *Drosophila* wing discs, while in vertebrates, collagen I mediates force propagation during neural crest and endothelial migration (Pastor-Pareja and Xu, 2011; Tang, 2020). Notably, many of our identified collagens are annotated as cuticle collagens, yet our findings reveal novel roles for them in internal ECM contexts. In addition, we identified DPY-18/P4HA1/2 and PDI-2/P4HB, enzymes required for collagen hydroxylation and disulfide-bond formation, as key regulators of gonad shape. Disrupting collagen processing compromises tissue mechanics, underscoring the broader principle that both collagen structure and post-translational modification are essential for morphogenetic control (Rahimi et al., 2022).

Previous work showed that VAB-3-dependent switching between the α-integrins INA-1 and PAT-2 is essential for gonad formation (Meighan and Schwarzbauer, 2007). INA-1, likely a laminin-binding α-integrin, supports early-stage migration, whereas PAT-2 is required for dorsal pathfinding and final positioning. PAT-2 is predicted to bind RGD motifs, which are found in some adult-enriched collagens such as COL-120 (Jayadev et al., 2019). However, many of the collagens critical for migration cessation lack canonical RGD motifs, suggesting functions independent of classical integrin–RGD binding. We propose two non-exclusive mechanisms: (1) the DTC may lay down a collagen matrix that it cannot degrade due to decreased expression of metalloproteases or incompatible protease profiles; and (2) non-RGD-mediated interactions between the collagen matrix and the DTC may trigger downstream changes in protease secretion, cytoskeletal remodeling, or polarity. Such pathways could convert the DTC into a more anchored, less migratory state. Selective integrin activation has been documented in other *C. elegans* tissues, such as the pharynx, where ECM incorporation occurs even without laminin, suggesting that integrin switching could reflect a broader ECM architectural shift from laminin-to collagen-rich environments (Jayadev et al., 2019; Jayadev et al., 2022).

An important open question is how the timing of collagen expression is coordinated with migration cessation. Type IV collagen α-chain ratios vary across tissues and stages, with LET-2:LET-2:EMB-9 trimers dominating the gonadal basement membrane (Srinivasan et al., 2025). This raises the possibility that compositional shifts in collagen architecture contribute to migration arrest. The heterochronic transcription factor LIN-29 is a strong candidate regulator, as it initiates the larval-to-adult transition and induces adult-specific collagen expression in the hypodermis (Rougvie and Ambros, 1995).

More recent studies show LIN-29 controls a broader suite of adult-enriched collagens and may coordinate ECM remodeling with developmental timing (Abete-Luzi et al., 2020). Additionally, microRNAs such as *lin-4* and the *let-7* family help synchronize cessation by repressing early regulators and enabling activation of adult-enriched collagens (Grishok et al., 2001; Zinovyeva et al., 2014). Together, these findings suggest that DTC migration arrest results from the integration of stage-specific transcriptional programs, ECM composition changes, and receptor–ligand interactions, providing a framework for understanding how mechanical and molecular cues converge to control tissue architecture.

In summary, we show that stage-specific collagen expression contributes to DTC migration cessation and gonad morphology in *C. elegans*. Our findings identify specific collagens required for this process and suggest that their regulation, mechanical properties, and potential signaling functions play a direct role in controlling tissue structure during development. More broadly, our findings underscore the general role of collagen in tissue morphogenesis and highlight conserved matrix features that may influence migration and tissue structure in other systems.

## Experimental Procedures

### Strains and Culture Conditions

*C. elegans* strains were maintained on Nematode Growth Medium (NGM) plates seeded with *E. coli* OP50 at 20°C under standard conditions (Brenner, 1974). The strain NK3026 (*let-2::mNG*) was used for collagen localization imaging. The RNAi-sensitized strain RZB353 (*cpIs121 I; rrf-3(pk1426) II; rde-1(ne219) V; ltIs44 [Ppie-1::mCherry::PH(PLC1delta1); unc-119(+)] IV*) was used for RNAi knockdown experiments. The strain NF2168 (*mig-24p::Venus (tkIs11)*) was used for fluorescence-activated cell sorting (FACS) and RNA-seq dataset generation, as it expresses Venus in the distal tip cells under control of the *hlh-12 (aka mig-24)* promoter.

### DTC collection and RNA sequencing

Distal tip cells (DTCs) were isolated from *Caenorhabditis elegans* at four developmental stages: late L3, early L4, late L4, and young adult as described in (Agarwal et al., 2025). The strain used for isolation was NF2168 [*mig-24p::Venus (tkIs11)*] (Tamai and Nishiwaki, 2007), which expresses a transcriptional GFP reporter driven by the *hlh-12* promoter, enabling specific labeling of DTCs. Three biological replicates were collected for each condition. cDNA was synthesized as described in (Agarwal et al., 2025). Briefly, sequencing libraries were prepared using the Nextera XT DNA Library Preparation Kit (Illumina) and sequenced using Illumina NextSeq500 with paired-end 2×150 □ bp reads. Library quality was assessed using FastQC (Andrews, 2010). Clean reads were aligned to the *C. elegans* ce11 reference genome using Bowtie 2 (Langmead and Salzberg, 2012). Gene-level read counts were generated using HTSeq-count (Anders et al., 2015) with the Ensembl release 94 GFF annotation.

### Normalization and Differential Expression Analysis

Gene count matrices were processed using the edgeR package in R. Genes with low expression were removed using the filterByExpr() function prior to normalization. Trimmed mean of M-values (TMM) normalization was then applied to account for compositional differences between libraries. Differential expression analysis was conducted using the glmQLFit() and glmQLFTest() functions. A design matrix was constructed to compare pooled larval DTCs (GL2–GL4) versus adult DTCs (GA), and also to compare GFP-positive versus non-GFP control samples at each stage. Significant genes were identified using Benjamini-Hochberg adjusted FDR < 0.05 and |log2 fold change| > 1 thresholds.

Normalized counts were also used to generate heatmaps and principal component analyses (PCA) across stages, enabling visualization of global trends and stage-specific expression dynamics.

### Enrichment and Category Analysis

WormCat was used to assign functional categories to differentially expressed genes (Holdorf et al., 2020). Genes were annotated based on Category 1 and Category 2 terms, which represent nested levels of functional classification: Category 1 encompasses broad biological processes (e.g., “Signaling,” “Transcription,” “Extracellular Material”), while Category 2 provides more specific sub-functions within each category (e.g., “Extracellular Material: Collagen,” “Signaling: G protein”). Enrichment scores and average log2 fold change were visualized as bubble plots. Collagen genes were extracted from the “Extracellular Material: Collagen” category for downstream analysis.

### Heatmaps and PCA

Normalized expression values were visualized using Morpheus (Broad Institute) and R-based heatmaps. Z-score normalization was applied by row to highlight relative expression dynamics. PCA was performed using the prcomp() function in R on TMM-normalized log2 CPM values.

### Clustering Analysis

Mfuzz was used to cluster collagen genes with significant differential expression across developmental stages. Prior to clustering, TMM-normalized counts per million (CPM) were z-score standardized across the four green-labeled conditions (GL2, GL3, GL4, GA) to emphasize temporal expression patterns independent of absolute magnitude. Clustering was performed on the standardized data using a fuzzifier parameter (m) estimated from the dataset. The optimal number of clusters was determined using the minimum centroid distance (Dmin) plot. For visualization and interpretability, average expression plots reflect TMM-normalized CPM values, while heatmaps are based on z-score standardized expression values to highlight relative differences across stages. For additional visualization in Morpheus, hierarchical clustering was applied to filtered collagen genes using Pearson correlation distance and average linkage to reorder gene rows in heatmaps.

### Volcano Plots and Gene Subsets

Volcano plots were generated in R using plotly for global DEGs, DTC migration genes, and collagens. TMM-normalized counts were used for DE testing, and significant genes were colored based on fold change and adjusted p-value. Plots were generated for pooled larval (GL2–GL4) versus adult (GA) DTCs.

### Expression Barplots and L2FC Comparisons

To compare collagen expression levels across developmental stages, TMM-normalized log_2_ counts per million (CPM) values were calculated using the edgeR package and averaged across replicates for each GFP-positive condition. Differential expression analysis was performed between GFP-positive and stage-matched GFP-negative samples using a generalized linear model (GLM) framework with quasi-likelihood F-tests to account for biological variability.

Resulting log_2_ fold changes (L2FC) were scaled and visualized in clustered heatmaps to assess differential expression of collagens and housekeeping genes. In addition, gene-wise barplots were generated for selected targets to highlight stage-specific enrichment patterns. All code and scripts used for this analysis are available at: **https://github.com/CramWormLab/DTC_Collagen_Paper**.

### Functional analysis

RNAi knockdown was performed by bacterial feeding using *E. coli* HT115 strains expressing dsRNA from either the Ahringer or Vidal RNAi libraries, or from custom constructs when necessary. Constructs were expressed from the L4440 vector. The strain used for DTC-specific knockdown was RZB353: *cpIs121 I; rrf-3(pk1426) II; rde-1(ne219) V; ltIs44 [Ppie-1::mCherry::PH(PLC1delta1); unc-119(+)] IV*. A minimum of 60 DTCs were scored per RNAi condition. Worms were plated on RNAi bacteria and imaged at 96 hours post-plating. RNAi plates contained 1 mM IPTG and 50 µg/mL ampicillin; 175 µL of induced bacterial culture was seeded per plate and dried overnight at room temperature. Worms were synchronized by bleaching, and eggs were plated directly onto RNAi plates. Worms were maintained at 20°C and transferred to fresh RNAi plates at 72 hours post plating to avoid starvation. In cases where target genes were not present in the Ahringer or Vidal RNAi libraries, gene-specific sequences were cloned by TA cloning into the L4440 backbone and insertion was confirmed by Sanger sequencing. The empty L4440 vector was used as a negative control. *tln-1* RNAi served as a positive control based on its known role in DTC migration (Cram et al., 2003). Importantly, the lethality of *tln-1* RNAi and other genes in standard RNAi backgrounds suggests that the observed phenotypes reflect DTC-specific knockdown in our sensitized strain.

DTC migration phenotypes were categorized into five groups: (A) wild-type, (B) overshoot, (C) swirl, (D) incorrect turn, and (E) other abnormal morphologies. Gonad width was measured 25 µm proximal to the DTC using the FIJI line tool, and worms were classified as having normal or abnormal morphology based on these measurements, which were conducted in parallel with migration scoring. Imaging conditions, strain backgrounds, and reporter usage were consistent with those described for DTC migration analysis. For gonad morphology assessment, at least 40 worms per condition were imaged, and each gene was tested in at least two independent experiments using freshly prepared plates and bacterial cultures. Results from multiple experiments were pooled. Gonad width comparisons between RNAi and control conditions were assessed using Welch’s *t*-test, and categorical phenotype distributions were evaluated with Fisher’s exact test. In both cases, Benjamini-Hochberg correction was applied to adjust for multiple testing. Quantification plots were generated using GraphPad Prism.

### Imaging and Microscopy

All DIC and fluorescence imaging was performed using a Nikon 80i microscope equipped with a 60x oil immersion objective. *let-2::mNG* transgenic worms (strain NK3026) were imaged at late L3, early L4, late L4, and adult stages (Soh et al., 2025). Images were processed using FIJI, and scale bars of 20 µm were included in all representative figures.

### Phylogenetic and Ortholog Analysis

Human orthologs for *C. elegans* collagen genes were identified based on sequence similarity. Protein sequences of collagens producing RNAi phenotypes were aligned using MAFFT. Phylogenetic trees were constructed using IQ-TREE with the LG+F+G4 model and 1000 bootstrap replicates. The gene *col-17* was excluded from the alignment due to extreme branch length. Trees were visualized using iTOL.

### STRING Network and Cytoscape Visualization

STRING networks were generated using the stringApp v2.2.0 plugin in Cytoscape (Cytoscape v3.8) with a confidence score cutoff of 0.7. Networks were visualized using the Prefuse Force Directed Layout with default heuristic weight interpretation (Doncheva et al., 2019; Shannon et al., 2003; Szklarczyk et al., 2021). Nodes representing genes without interactions (singlets) were excluded. Node color reflected the developmental stage at which each gene exhibited peak expression (Late L3, Early L4, Late L4, or Adult), based on normalized TMM expression values. Tightly clustered subregions, such as the adult-enriched collagen group, were highlighted based on their natural network structure and not algorithmic clustering.

### Figure Preparation

All figures were prepared using R, Morpheus, and finalized in Adobe Photoshop. Volcano plots, heatmaps, clustering plots, PCA plots, and STRING networks were generated in RStudio. Bar graphs and morphology dot plots were created using GraphPad Prism. All non-fluorescent images were adjusted for optimal brightness and contrast to enhance visual clarity.

## Acknowledgements

We thank members of the Cram, Apfeld, and Vollmer labs for helpful discussions, and to the Sherwood lab for helpful insights into *C. elegans* collagens. We are especially grateful to Sophia Guerra for assistance with figure illustration and Luke Goncalves for helpful technical support. Some strains were provided by the CGC, which is funded by NIH Office of Research Infrastructure Programs (P40 OD010440). This work was supported by a grant from the Binational Science Foundation to R.Z.B (2021014).

