## Supplemental Tables 1 and 2 for "A switch in collagen expression regulates cessation of distal tip cell migration in *C. elegans*"

**Supplementary Table 1.** List of genes screened in this study. Genes are organized by sequence name, locus, functional class, and associated distal tip cell (DTC) migration phenotype. Values indicate frequency of observed defects and number of animals scored.

**Supplementary Table 2.** Human orthologs of collagen genes. *C. elegans* collagen genes identified in this study are paired with their closest human orthologs, highlighting conservation of extracellular matrix function.

**Supplementary Table 1: List of genes screened in this study**

| Sequence Name | Locus | Functional Class | P0 defect | n |
| --- | --- | --- | --- | --- |
| Y71G12B.11 | *tln-1* | Cytoskeleton | 0.99 | 163 |
| C09G5.3 | *col-79* | Extracellular material | 0.03 | 37 |
| T07H6.3 | *col-166* | Extracellular material | 0.05 | 41 |
| F57B7.3 | *col-156* | Extracellular material | 0 | 44 |
| T11B7.3 | *col-118* | Extracellular material | 0.26 | 61 |
| F15A2.1 | *col-178* | Extracellular material | 0.19 | 62 |
| C27H5.5 | *col-36* | Extracellular material | 0.02 | 55 |
| W08D2.6 | *col-123* | Extracellular material | 0 | 56 |
| Y11D7A.11 | *col-120* | Extracellular material | 0.04 | 57 |
| C46A5.3 | *col-14* | Extracellular material | 0.02 | 57 |
| W03G11.1 | *col-181* | Extracellular material | 0.02 | 57 |
| F54C9.4 | *col-38* | Extracellular material | 0.13 | 60 |
| F11H8.3 | *col-8* | Extracellular material | 0.22 | 60 |
| M199.5 | *col-135* | Extracellular material | 0.02 | 61 |
| K02D7.3 | *col-139* | Extracellular material | 0.11 | 62 |
| Y73B6BL.34 | *dpy-18* | Extracellular material | 0.25 | 63 |
| F56B3.1 | *col-103* | Extracellular material | 0.15 | 67 |
| F29C4.8 | *col-99* | Extracellular material | 0.04 | 67 |
| M18.1 | *col-129* | Extracellular material | 0.07 | 68 |
| F38A3.1 | *col-81* | Extracellular material | 0.03 | 68 |
| F41F3.4 | *col-184* | Extracellular material | 0.11 | 69 |
| C34F6.2 | *col-93* | Extracellular material | 0.12 | 69 |
| F11G11.11 | *col-20* | Extracellular material | 0.06 | 70 |
| F59E12.12 | *bli-2* | Extracellular material | 0.04 | 72 |
| C24F3.6 | *col-124* | Extracellular material | 0.04 | 72 |
| F55C10.3 | *col-155* | Extracellular material | 0.14 | 73 |
| F11G11.10 | *col-17* | Extracellular material | 0.47 | 73 |
| C53B4.5 | *col-119* | Extracellular material | 0.19 | 74 |
| Y69H2.14 | *bli-6* | Extracellular material | 0.11 | 76 |
| T01B7.7 | *rol-6* | Extracellular material | 0.09 | 76 |
| ZK1193.1 | *col-19* | Extracellular material | 0.13 | 78 |
| F30B5.1 | *dpy-13* | Extracellular material | 0.65 | 79 |
| W05B2.5 | *col-140* | Extracellular material | 0.11 | 82 |
| F26F12.1 | *col-42* | Extracellular material | 0.11 | 84 |
| F43G9.11 | *col-101* | Extracellular material | 0.17 | 95 |
| T15B7.3 | *col-143* | Extracellular material | 0.18 | 96 |
| C15A11.5 | *col-7* | Extracellular material | 0.13 | 117 |
| T06E4.6 | *col-146* | Extracellular material | 0.13 | 119 |
| ZK265.2 | *col-63* | Extracellular material | 0.11 | 157 |
| C07A12.4 | *pdi-2* | Protein modification | 0.16 | 63 |

**Supplementary Table 2: Human orthologs of collagen genes**

| *C. elegans* gene | Human Ortholog |
| --- | --- |
| *bli-1* | collagen type XXVII alpha 1 chain and collagen type IX alpha 1 chain. |
| *cle-1* | collagen type XV alpha 1 chain and collagen type XVIII alpha 1 chain. |
| *col-119* | collagen type IV alpha 2 chain. |
| *col-123* | collagen type VI alpha 2 chain |
| *col-135* | collagen type IX alpha 2 chain. |
| *col-14* | collagen type IX alpha 1 chain. |
| *col-141* | collagen type VI alpha 2 chain. |
| *col-152* | collagen type XXI alpha 1 chain. |
| *col-175* | collagen type XXI alpha 1 chain. |
| *col-179* | collagen type IX alpha 2 chain |
| *col-181* | collagen type V alpha 3 chain and |
| *col-33* | collagen type XXI alpha 1 chain. |
| *col-36* | collagen type XXI alpha 1 chain. |
| *col-77* | collagen type VI alpha 2 chain. |
| *col-80* | collagen type XXI alpha 1 chain. |
| *col-99* | collagen type XXIII alpha 1 chain and collagen type XXV alpha 1 chain. |
| *cutl-23* | collagen type VI alpha 3 chain |
| *dpy-14* | collagen type III alpha 1 chain |
| *dpy-4* | collagen type XXVII alpha 1 chain, collagen type IV alpha 2 chain and collagen type IX alpha 1 chain. |
| *dpy-9* | collagen type IV alpha 5 chain |
| *let-2* | collagen type IV alpha 1 chain, collagen type IV alpha 5 chain and collagen type IV alpha 6 chain. |
| *rol-6* | collagen type III alpha 1 chain |
